# Measurements of morphological and biochemical alterations in individual neuron cells associated with early neurotoxic effects in Parkinson’s disease

**DOI:** 10.1101/080937

**Authors:** Su-A Yang, Jonghee Yoon, Kyoohyun Kim, YongKeun Park

## Abstract

Parkinson’s disease (PD) is a common neurodegenerative disease. However, therapeutic methods of PD are still limited due to complex pathophysiology in PD. Here, we present optical measurements of individual neurons from *in vitro* PD model using optical diffraction tomography (ODT). By measuring 3-D refractive index distribution of neurons, morphological and biochemical alterations in *in-vitro* PD model are quantitatively investigated. We found that neurons show apoptotic features in early PD progression. The present approach will open up new opportunities for quantitative investigation of the pathophysiology of various neurodegenerative diseases.

---

Parkinson’s disease (PD) is a prevalent neurodegenerative disease that is distinctly marked by loss of control of motor functions in patients^1^. Clinical symptoms of PD involve tremor at rest, rigidity or slowness of voluntary movements, and postural instability. These symptoms are known to be caused by progressive degeneration of dopaminergic neurons located in the substantia nigra. Previous reports showed that the genetic defect in complex I of the mitochondrial electron transport chain produces reactive oxygen species in dopaminergic neurons which result in neurodegeneration^2^. Although the pathophysiology of PD has been intensively studied using *in vitro* and *in vivo* models, molecular mechanisms underlying neurotoxicity in PD remain unclear, and thus therapeutic methods are limited. Moreover, there are little methods to detect early neurodegeneration in PD.

To measure neurodegenerative progress in neurons, various optical methods are intensively used. Light microscopy, such as bright-field microscopy or phase contrast microscopy, which is a basic tool for *in vitro* study, conveniently visualizes cellular morphology^3, 4^, but it provides only qualitative information. On the other hand, confocal and multi-photon fluorescence microscopy allows acquisition of three-dimensional (3-D) structural and molecular images of neurons in *in vitro* and *in vivo* models with a high spatial resolution^5, 6^. However, microscopy which involves the use of fluorescence requires invasive labeling procedures such as chemical staining or genetic modification, which inevitably induces phototoxicity or photobleaching. Lastly, atomic force microscopy (AFM) has been used to measure cellular morphology, for example, the cellular height in *in vitro* neurodegeneration model. AFM provides quantitative structural information of neurons with immensely high resolution, but it requires long operation time due to scanning of an AFM tip to measure the whole cellular area^7^.

To overcome aforementioned limitations and quantitatively measure early neurodegenerative progress in the PD model, quantitative phase imaging (QPI) techniques^8, 9^ can be utilized. Based on the principle of holography, QPI techniques provides quantitative and label-free images of live cells and tissues, exploiting the refractive index (RI) distribution of optical contrasts. QPI techniques have been applied to study the pathophysiology of various biological samples, including haematology^10^, malaria infection^11, 12^, sickle cell diseases^13^–^16^, and microbiology^17, 18^. In particular, 2-D QPI techniques or digital holographic microscopy techniques had been utilized for the study of biophysics of neuron cells^19, 20^ and for imaging alterations in brain tissue slices caused by Alzheimer’s disease^21^.

Here, we report the investigation of early neurotoxic effects in an *in vitro* PD model using 3-D QPI techniques. Exploiting optical diffraction tomography (ODT)^22^–^24^, 3-D RI tomograms of samples are measured at an individual cell level, which provides quantitative morphological and biochemical information. Because RI is an intrinsic optical property of materials, ODT does not require any invasive labeling process. Recently, measuring 3-D RI distributions has been widely applied to study the pathophysiology of various biological samples, such as red blood cells^25-28^, white blood cells^29, 30^, cancer cells^31^–^37^, phytoplankton^38^, and bacteria^39^.

Here, we report the investigation of early neurotoxic effects in an *in vitro* PD model using ODT. To establish the *in vitro* PD model, we used 1-methyl-4-phenylpyridinium ion (MPP+) which is a metabolite of 1-methyl-4- pheneyl-1,2,3,6-tetrahydropyridine (MPTP) that is known to cause symptoms of PD^40, 41^. For studying the early pathophysiology of PD, we performed optical measurements on the morphological and biochemical properties of the cells after 5 hours of MPP+ treatment. The quantitative characteristics of the cells, including cellular surface area, volume, sphericity, height, dry mass density, and dry mass, are retrieved from measured 3-D RI distributions. We found that early neurotoxic effects of MPP+ change the cellular morphology of MPP+-treated cells to become round and shrinkage while cellular dry mass remains the same, which are well-known apoptotic features. The demonstration of ODT in the neurodegenerative disease research will provide new opportunities for quantitative investigation of pathophysiology in PD as well as other neurodegenerative diseases such as Alzheimer’s disease and amyotrophic lateral sclerosis.

## Results

### Label-free 3-D quantitative measurements of SH-SY5Y cells using optical diffraction tomography

To perform quantitative imaging and analysis of neurons at an individual cell level, we measured 3-D RI distributions of the cells using 3-D quantitative phase imaging techniques. A Mach-Zehnder interferometry equipped with a dual-axis galvanomirror [Fig. 1(a)], which is comparable to the setup used in previous work^28, 42^. Human neuroblastoma dopaminergic cells, SH-SY5Y, were used in this study. A SH-SY5Y cell was illuminated with a coherent laser beam. Multiple holograms of the SH-SY5Y cell were measured by changing angle of illumination using a dual-axis galvanomirror, and then optical field information (amplitude and phase) of the sample were retrieved from measured holograms via an optical field retrieval algorithm [Fig. 1(b)]. From the measured multiple 2-D amplitude and phase information, a 3-D RI tomogram was reconstructed via optical diffraction tomography algorithm (See Method).

**Figure 1.**
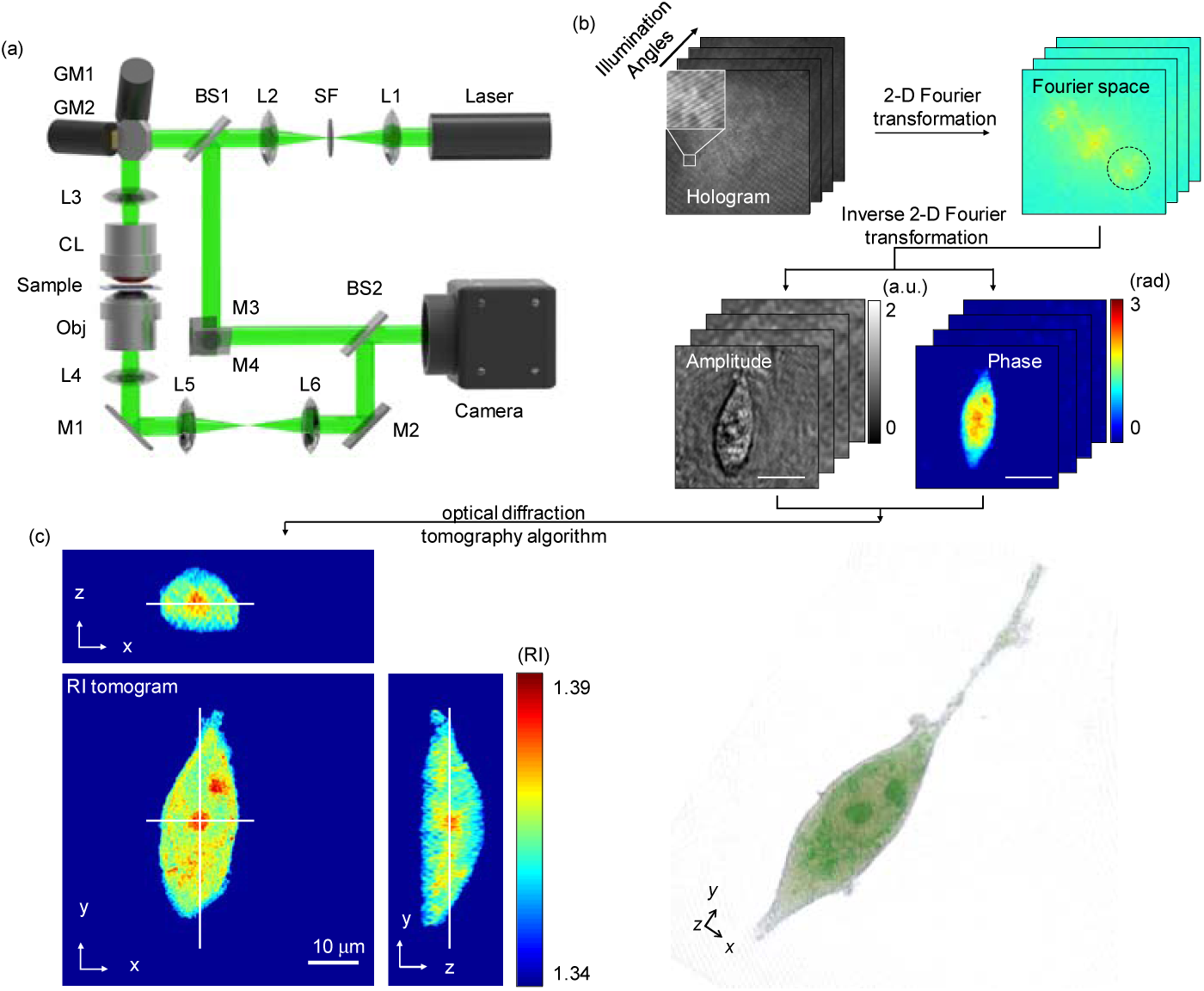
Schematics of the experimental setup and procedures for measuring 3-D refractive index tomogram of a SH-SY5Y cell. (a) A Mach-Zehnder interferometric microscope equipped with a 2-D scanning galvanomirror (GM). BS1−2, beam splitters; L1−6, lenses; SF, spatial filter; CL, condenser lens; OL, objective lens; M1−4, mirrors. (b) Field retrieval procedures from holograms measured with various angles of illumination (top, left). Inset: enlarged view of spatially modulated interference patterns. From the 2-D Fourier spectrum (top, right), the amplitude and phase information are retrieved (bottom). Scale bar is 10 μm. (c) Cross-sectional slices of a RI tomogram of a SH-SY5Y cell. The RI tomogram is reconstructed from the multiple 2-D amplitude and phase information via optical diffraction tomography algorithm. (d) A 3-D rendered image of (c) using customized transfer function.

Figure 1(c) shows cross-sectional slices of a reconstructed tomogram of a SH-SY5Y cell on various axes. The 3-D RI distribution of the cell clearly shows cell boundaries and intracellular organelles such as nucleus and nucleoli, based on their distinct distributions of refractive index values. For the visualization purpose, the 3-D RI tomogram was rendered using commercial software (Tomostudio, Tomocube Inc., Republic of Korea), as shown in Fig. 1(d) and Supplementary Movie 1. The 3-D rendered image shows the entire RI distribution of the cell, which presents cell morphology including cell body, axon, and intracellular components.

### Quantitative morphological and biochemical analysis of early neurodegenerative effects using the in vitro PD model

To systematically investigate early pathophysiological features of PD in individual SH-SY5Y cells, quantitative morphological and biochemical characteristics of the control group and PD model are analysed. From the measured 3-D RI distributions, quantitative morphological (i.e. surface area, cellular volume, sphericity, and cellular height) information of the cells can be calculated (See Methods).

Figure 2 shows results of quantitative characterizations of the control and MPP+-treated group. The mean values of cellular surface areas of individual SH-SY5Y cells are 1,290.52 ±339.67 and 1,106.95 ±321.25 μm^2^, for the control and MPP+-treated group, respectively [Fig. 2(a)]. There is a statistical difference (*p*-value < 0.01) of cellular surface areas between the two groups. The mean values of cellular volumes of two groups are 1371.74 ± 488.84 and 1415.32 ±504.71 fL, respectively [Fig. 2(b)]. From the calculated cellular surface areas and volumes, sphericity is directly obtained. The calculated mean sphericity values of the control and MPP+-treated group are 0.47 ±0.07 and 0.56 ±0.09, respectively [Fig. 2(c)]. In comparison to the control group, the MPP+-treated group shows a statistical difference (*p*-value < 0.01) in sphericity values, which indicates that MPP+-treated cells had become round. The cellular height information also can be retrieved from 3-D RI tomograms. The mean values of cellular height are 10.23 ±1.31 and 11.73 ±1.66 μm for the control and MPP+-treated group, respectively [Fig. 2(d)]. There is a statistical difference (*p*-value < 0.01) in cellular height between the two groups. Therefore, it was observed that SH-SY5Y cells became thicker and more round in shape upon MPP+ treatment, while cellular volume remained static.

**Figure 2.**
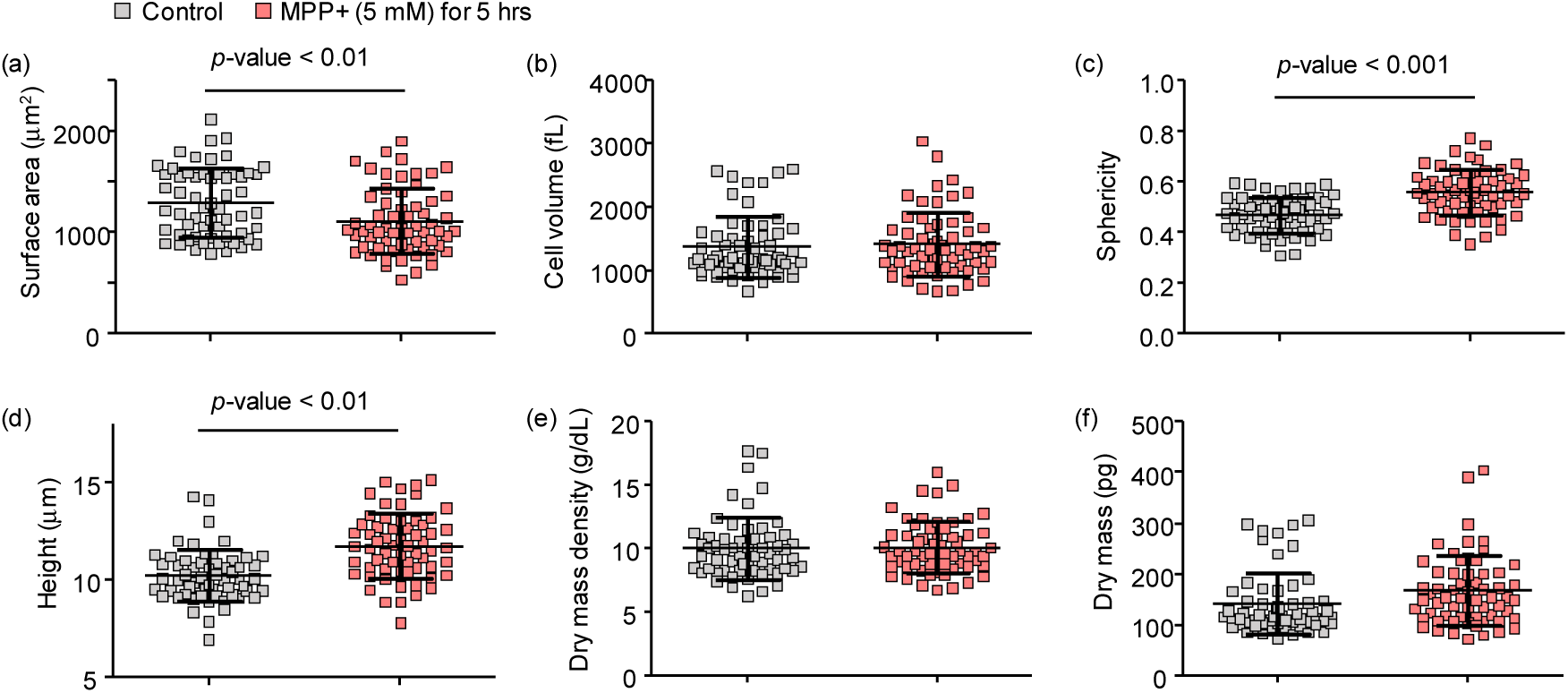
Quantitative analysis of morphological and biochemical information of control (*n* = 59) and MPP+ treated group (*n* = 60). (a) Surface area, (b) cell volume, (c) sphericity, (d) height, (e) dry mass density, and (f) dry mass. Each symbol indicates an individual cell measurement and the black horizontal line indicates the mean value with the vertical line of standards deviation. *p*-values are calculated by Student’s *t*-test.

Next, we calculated biochemical information (i.e. cellular dry mass density and dry mass) of individual cells to test whether MPP+ treatments affect intracellular protein contents. RI values can be converted to local dry mass density because of their linear relationship to protein concentration (See Methods). Total cellular dry mass is then obtained by integrating the dry mass density over the cellular volume. The calculated mean values of dry mass density are 10.00 ±2.43 and 10.09 ±2.02 g/dL, and the mean values of dry mass are 141.75 ±61.68 and 166.86 ±68.16 pg for the control and MPP+-treated group, respectively [Figs. 2(e-f)]. These results indicate that 5 hours of MPP+ treatments do not cause statistical differences in intracellular protein contents of MPP+ treated cells in comparison to the untreated cells. Therefore, we found that early effects of MPP+ treatment on SH-SY5Y cells involve alterations of the cellular surface area and thickness, rather than changes of the cellular volume and intracellular protein contents. These morphological changes are consistent with the major hallmarks of a cell undergoing apoptosis^43, 44^.

### Discrimination of SH-SY5Y cells in healthy, apoptotic and mitotic states

In order to closely examine the features of SH-SY5Y cells upon MPP+ treatment, we performed analysis of MPP+ effects at an individual cell level. Figure 3 shows three representative SH-SY5Y cells in different cellular condition: healthy, apoptotic, and mitotic states. Figure 3(a) shows a 3-D rendered image of a healthy SH-SY5Y cell that has elongated morphology with a nucleus and an axon. In contrast, the apoptotic cell exhibits cell shrinkage, membrane bleb formations and condensed intracellular components, which are well-known apoptotic phenotypes [Fig. 3(b)]^43, 44^. Moreover, the quantitative analysis reveals that the sphericity and cellular height of the apoptotic cell are higher than that of the healthy cell, which indicates that apoptotic cell’s morphology has been altered to a rounder shape. We also observed SH-SY5Y cells in mitotic states. Figure 3(c) shows a representative 3-D rendered image of a mitotic cell and its quantitative characteristics. The mitotic SH-SY5Y cell clearly shows two nucleus and nucleoli. Even though cellular morphology of the mitotic cell is similar to that of the healthy cell, the total dry mass of the mitotic cell is two times higher than that of the healthy cell. This result is consistent with previous reports that have demonstrated that the cellular mass increases twice during mitosis as it produces intracellular proteins in preparation for splitting into two daughter cells^45^. In addition, the mitotic cell has higher sphericity value compared to the healthy cell because cells in mitosis become round prior to cell division. Thus, the mitotic cell and apoptotic cell have similar sphericity values due to their round shapes^46^.

**Figure 3.**
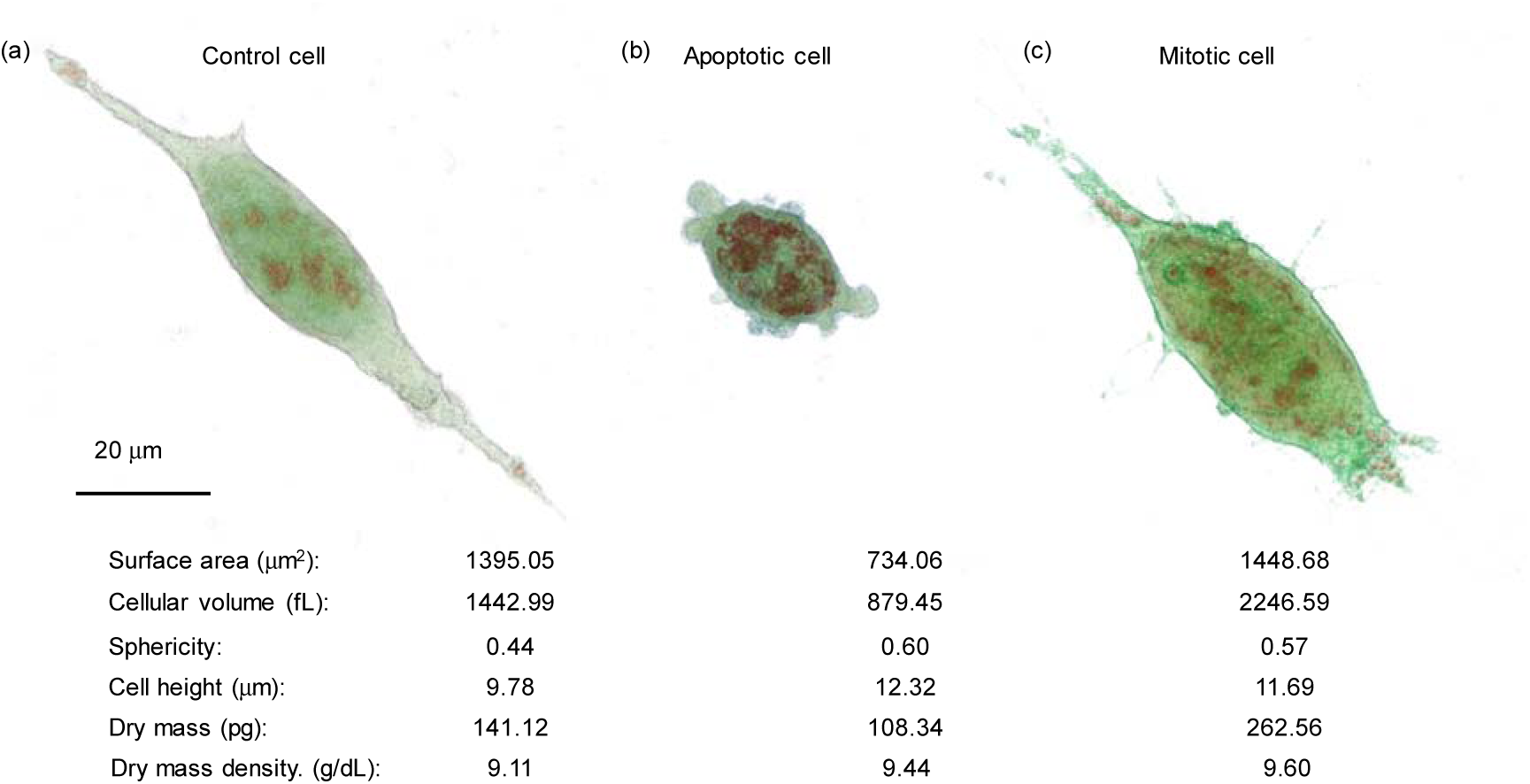
Representative 3-D rendered tomograms and quantitative characterization of SH-SY5Y cells in different conditions. (a) Control cell, (b) apoptotic cell, and (c) mitotic cell. Scale bar is 20 μm.

To discriminate SH-SY5Y cells according to their cellular condition, we plotted sphericity values over the cellular dry mass of SH-SY5Y cells with and without MPP+ treatment. Figure 4 shows the 2-D scatter plot with and without MPP+ treatments and clustering of cells according to cellular states: health, apoptotic, and mitotic states. The cells are classified to three cellular states using a *K*-means clustering algorithm^47^. The blue, green, and red colours in Fig. 4 indicate cellular states such as healthy (*n* = 56), apoptotic (*n* = 41), and mitotic stages (*n* = 22), respectively. The healthy cells (low SI values, low dry mass) are localized on the lower left quadrant of the 2-D plot of which the mean sphericity value and cellular dry mass are 0.43 ±0.05 and 122.25 ±29.48 pg, respectively. The apoptotic cells (high SI values, low dry mass), located above the healthy cells, exhibit higher sphericity (0.60 ±0.06) and similar dry mass (134.27 ±31.67 pg) compared to the healthy cells. On the other hand, the mitotic cells (high SI values, high dry mass) exhibit increases in both sphericity values (0.54 ±0.04) and dry mass (252.83 ±32.78 pg) in comparison to healthy cells, thus they are localized on the right side of the plot. We observed that MPP+-treated cells exhibit a high ratio of apoptosis (30/60, 50%) compared to the control group (11/59, 18.6%). These results show that the MPP+ treated group has a tendency toward the apoptotic, suggesting the MPP+ causes neurodegenerative effects in SH-SY5Y cells and induces apoptosis within 5 hours of the treatment.

**Figure 4.**
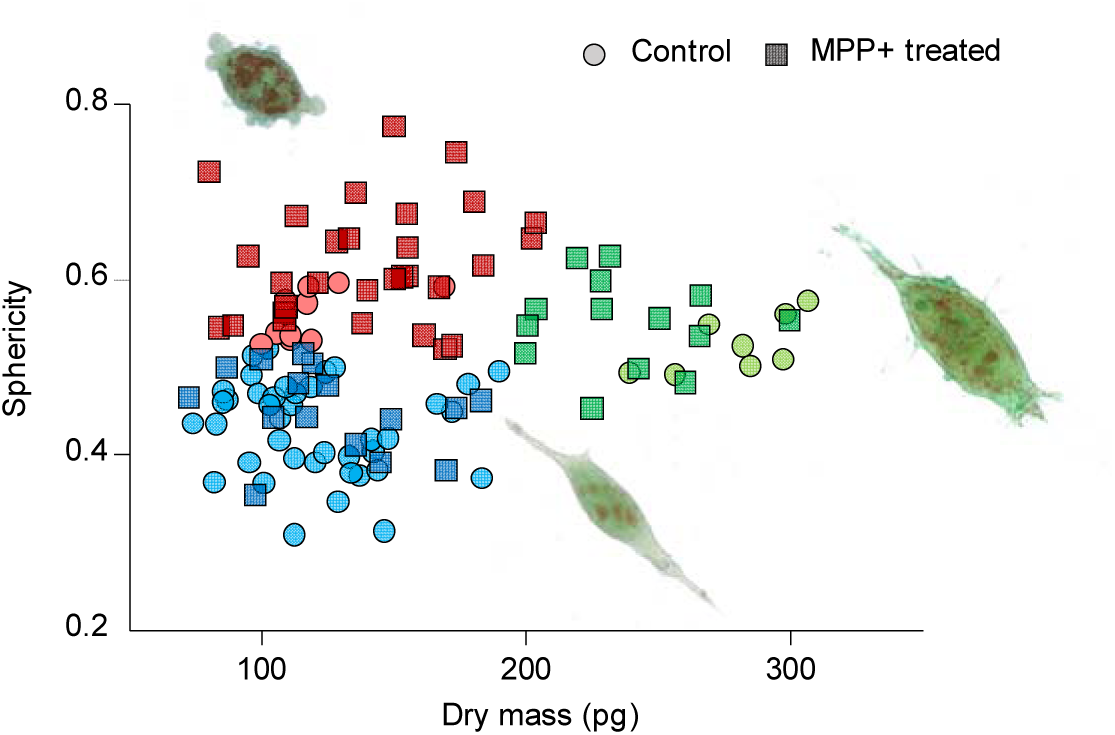
2-D plot of sphericity over dry mass. *K*-means clustering of control and MPP+ treated group using cellular dry mass and sphericity. The circle dots and square dots represent control and MPP+ treated groups, respectively. The blue, green and red colours indicate cellular status such as healthy, mitotic, and apoptotic cells, respectively. 3-D rendered tomograms near the dots show the representative morphology of each cellular status.

### Discussion

In summary, we measured 3-D RI tomograms of neurons using a 3-D holographic microscope and demonstrated that quantitative morphological and biochemical information obtained from RI distributions of measured cells enable investigation of the early pathophysiology of Parkinson’s disease. From quantitative analysis of cellular morphology, we observed that MPP treatments alter cellular shapes from flat to round shape within 5 hours of MPP+ treatment, while cellular dry mass remains static. The result was shown to be consistent with the previous observations of apoptotic features of a neuronal cell affected by MPP+. Furthermore, using the 2-D plot of sphericity over cellular dry mass, we found that healthy, mitotic or apoptotic status of a cell can be discriminated in an MPP+ treated population. Thus, our method provides an effect tool to report whether an individual cell has been affected by MPP+ and is undergoing apoptosis.

Our results from quantitative morphological and biochemical analysis of SH-SY5Y cells indicated that MPP+ causes apoptosis rather than necrosis. The MPP+ treatment alters cellular morphology to become rounder and form membrane blebs, which are typical apoptotic morphology^43, 44^. These morphological changes are consistent with previous reports that MPP+ induces disruption of the cytoskeleton in neurons^48, 49^. Furthermore, we confirmed that membrane integrity of the MPP+-induced apoptotic cells remains stable by showing cellular dry mass are similar regardless of MPP+ treatments; since if there is a loss of membrane integrity of a cell, the cellular dry mass would be rapidly decreased by leaking intracellular components to extracellular regions. Thex results are consistent with the previous studies, which observed that MPP+ causes apoptosis in SH-SY5Y cells, featuring membrane blebs, cytoskeleton disruption, neurite shortening, cell shrinkage and rounding.^40, 48, 50–52^ Therefore, we conclude that SH-SY5Y cells undergo apoptosis within 5 hours of MPP+ treatment.

To our knowledge, we are the first to use ODT to observe and report apoptotic features of MPP+ treated live SH-SY5Y cells. The ODT measures an RI distribution of the sample, which is an intrinsic optical property of material. Its advantage includes that it does not require any exogenous labeling procedures such as fluorescent staining or genetic modification that may cause disturbances in cellular function. Furthermore, elimination of such experimental steps has saved time and reduced the possibility of errors. Also, quantitative morphological and biochemical information enable discrimination of apoptotic cells from healthy and mitotic cells without using apoptosis marker such as annexin V, which stains phosphatidylserine on cellular plasma membrane^53, 54^. The present method possesses the unique ability to distinguish and investigate only the specific cell population that shows MPP+-induced neurotoxicity at an individual cell level. This provides more accurate pathophysiological information of PD than by analysing mixed cell population that includes the cells not affected by MPP+.

The limitation of the present method is a lack of molecular specificity. Unlike conventional biochemical methods, such as western blots and polymerase-chain reactions or fluorescent techniques, RI distribution does not provide specific molecule information because refractive index increments required to convert RI values to the local concentration of intracellular molecules are similar^55^. Only several intracellular components including lipid droplets, nucleus, and nucleoli can be identified due to their distinctive shapes and RI values^23^. Although the molecular specificity of the present label-free RI measurement technique may not be as high as that of labeling techniques or conventional biochemical methods, it could be improved by measuring spectroscopic RI distributions, which can separate molecules spectrally owing to their different optical dispersion properties^36^.

In conclusion, we envision that ODT will be useful in the study of pathophysiology in PD due its ability to investigate quantitative alterations of morphological and biochemical features of neurons in the disease condition. Because other neurodegenerative diseases such as Alzheimer’s disease or amyotrophic lateral sclerosis also cause abnormal changes in neuronal morphology and function, the present approach can be readily applied to study the pathophysiology of various neurodegenerative diseases.

## Methods

### SH-SY5Y cell preparation and creating Parkinson’s disease model

Human neuroblastoma, SH-SY5Y, cell lines were purchased from Korean Cell Line Bank. SH-SY5Y cells were cultured in Dulbecco’s modified Eagle’s medium with nutrient mixture F-12 (DMEM/F12, Welgene, South Korea) supplemented with 10% fetal bovine serum (Welgene, South Korea) and maintained at a 37°C, 5% CO_2_ incubator. Prior to 3-D holographic imaging, SH-SY5Y cells were detached from a culture flask using 0.25% Trypsin-EDTA (Welgene, South Korea) and placed on a 24 mm × 40 mm poly-L-lysine coated coverslip (Marienfeld-Superior, Germany) and maintained for 12 hours. Then, 5 mM MPP+ iodide (Sigma-Aldrich, USA)dissolved in autoclaved distilled water was treated to cells, and then, it was left in a dark environment for 5 hours before the experiment. The cells were washed with Dulbecco’s Phosphate-Buffered Saline (DPBS, Welgene, South Korea) immediately before the measurement. Next, 500 μL of pre-warmed DMEM/F12 was added to the coverslip, and then, another coverslip was placed on top of the cells to prevent the sample from drying.

### Optical diffraction tomography

In optical diffraction tomography, optical fields were measured by Mach-Zehnder interferometry [Fig. 1(a)]. A laser beam from a diode-pumped solid state laser (λ = 532 nm, 100 mW, Shanghai Dream Laser Co., Shanghai, China) is split into two arms using a beam splitter. One arm is used as a reference beam, and the other is used to illuminate the sample with changing angles using a dual-axis galvanomirror (GVS012, Thorlabs, Newton, NJ, USA). The sample is placed between a condenser lens (UPLSAPO Water 60×, numerical aperture (NA) = 1.2, Olympus, Japan), which was immersed in DPBS solution during imaging and an objective lens (PLAPON Oil 60×, NA = 1.2, Olympus, Japan). The diffracted beam from the sample was collected and interfered with the reference beam, generating spatially modulated interferograms that were recorded on a CMOS camera (1024 PCI, Photron USA Inc., San Diego, CA, USA). Typically, 300 holograms measured by different illumination angles were used for reconstructing a 3-D RI tomogram. From measured holograms, amplitude and phase information were retrieved via a field retrieval algorithm^56^. Then, a 3-D RI tomogram was reconstructed using retrieved optical fields information via the optical diffraction tomography algorithm. To fill missing information due to the limited NAs of the condenser and imaging objective lenses, a regularization algorithm based on non-negativity has been used^57^. Detailed information of ODT and the source code can be found elsewhere^25^. Recently, ODT systems have been commercialized by a few companies, including Holotomography (Tomocube Inc., Republic of Korea), which will facilitate the applications of the method presented in this study.

### Quantitative analysis

Quantitative morphological and biochemical information were calculated from reconstructed 3-D RI tomograms. To calculate all quantitative information of a sample, the voxels which have higher RI values than the background RI value were selected. From selected voxels, cell surface area *S* and volume *V* can be calculated corresponding to cell boundaries and the numbers of voxels, respectively. Then, sphericity is directly obtained from the measured surface area and volume, as follows: *Sphericity* = π^1/3^⋅(6*V*)^2/3^/*S*. The sphericity is a dimensionless parameter which indicates how round object is. The RI information also provides a local concentration of non-aqueous molecules including proteins, lipids, and nucleic acids inside cells^58, 59^. From the average RI value of the cells, dry mass concentration *ρ* was calculated. RI values of the cytoplasmic region of the cells are proportional to non-aqueous molecules (mostly proteins) as following relationship *n* = *n*_0_ + *αρ*, where *n* is RI of proteins, *n*_0_ is RI of surrounding media, and *α* is the refractive index increment (RII) of proteins. Because most proteins have similar RII values, we used the RII values of 0.2 mL/g in this study^60^. Total dry mass of a sample is calculated by simply integrating dry mass density over cellular volume.

### Statistical Analysis

P values are calculated by Student’s *t*-test comparing the cellular properties between various the normal control group and MPP+-treated cells. All the numbers follow the ±sign in the text is a standard deviation.

### Imaging processing

Image processing was performed using Matlab R2015b (Natick, MA, USA) and imageJ (NIH, USA) software. The 3-D image of the RI distribution of a sample is rendered by Tomostudio (Tomocube Inc., Korea).

## ACKNOWLEDGEMENTS

This work was supported by KAIST, and the National Research Foundation of Korea (2015R1A3A2066550, 2014K1A3A1A09063027, 2012-M3C1A1-048860, 2014M3C1A3052537) and Innopolis foundation (A2015DD126).

## AUTHOR CONTRIBUTIONS

Y.P. conceived the idea and directed the work. J.Y. and K.K. designed optics system and developed algorithms for experiments. S.Y. and J.Y established *in vitro* model and performed experiments. All authors wrote the manuscript.

